# Automated detection of macropods in Tasmania using drone surveys and convolutional neural networks (CNNs)

**DOI:** 10.1101/2025.11.26.688741

**Authors:** Yee Von Teo, Darren Turner, Jessie C. Buettel, Barry W. Brook

## Abstract

Manual annotation of drone imagery is labour-intensive and prone to observer bias, particularly when applied to large datasets across varied environments. To address this, a deep-learning pipeline was developed and evaluated for identifying macropods in visual (RGB) drone imagery, using a convolutional neural network (CNN) adapted from the `DeepForest` framework. The model was trained on annotated images of Forester kangaroos (*Macropus giganteus tasmaniensis*) and Bennett’s wallabies (*Notamacropus rufogriseus*) collected across two Tasmanian study sites. Performance was assessed using independent test sets from each site, representing open and forest-edge habitats, as well as a combined multi-site test set. Detection accuracy was quantified using precision, recall, and F1 scores, with further analyses evaluating the effect of solar altitude angle on model performance. The model achieved high recall across sites, indicating strong potential for minimising missed detections under diverse conditions. These results demonstrate the feasibility of applying transfer learning to drone-based wildlife surveys and highlight the promise of deep learning models for reducing manual effort in macropod monitoring, with applications for broader conservation and management workflows.

## 1. Introduction

Large mammalian herbivores have significant impacts on ecosystems (Huntly 1991; Bakker *et al*. 2006; Morgan 2021). They alter plant community diversity and ecosystem processes across a range of habitats (Huntly 1991; Rueda et al. 2013; Staver et al. 2021). Many large herbivores also act as ecosystem engineers, creating physical disturbances that generate microhabitats for other species and increase landscape heterogeneity (Wilby, Shachak & Boeken 2001). However, changes in their abundance can trigger cascading effects throughout food webs, ultimately altering biodiversity and ecosystem function (Wilby, Shachak & Boeken 2001). Intensive herbivory pressure can result in vegetation degradation through overgrazing or selective grazing/browsing, potentially causing shifts in plant community composition that favour less palatable or invasive species (Wardle 2001; O’Reilly-Wapstra & Cowan 2010).

In Australia, most large native herbivores belong to the family *Macropodidae*, which includes kangaroos, wallabies, and pademelons (collectively known as ‘macropods’) (Southwell *et al*. 1999; Chapman 2003). Through selective grazing on grasses and shrubs, macropods help maintain and shape vegetation structure, promote plant diversity, and maintain open habitats important for other species. However, when macropod populations reach high densities, overgrazing can occur, leading to the degradation of native plant communities, declines in biodiversity, and altered ecosystem dynamics (Ramsey & Wilson 1997; Wiggins *et al*. 2010; Gordon *et al*. 2021). During droughts or periods of limited forage availability, their feeding pressure may intensify, contributing to soil degradation and reduced pasture availability (Le Mar, Southwell & McArthur 2001).

On the island state of Tasmania, macropods are commonly found in agricultural landscapes where food is abundant, and predator pressure is low. In these areas, high local densities can result in significant browsing damage caused by native herbivores such as the Tasmanian pademelon (*Thylogale billardierii*) and Bennett’s wallaby (*Macropus rufogriseus*), reducing the growth of eucalypt seedlings in Tasmanian forestry plantations (Bulinski 2000). Their presence in agricultural settings has also led to conflicts with landholders due to the loss of pasture and crop production (Le Mar, Southwell & McArthur 2001). To manage growing macropod populations, the Tasmanian government has issued shooting permits for landholders to control the population of Tasmanian pademelons and Bennett’s wallabies on their properties (Wiggins *et al*. 2010). These species are harvested either recreationally or via property protection permits at a localised level. Since 1975, the Department of Natural Resources and Environment Tasmania (NRE Tas) has been carrying out annual nocturnal spotlight surveys to detect population trends of priority harvested species, including Tasmanian pademelons and Bennett’s wallabies, to ensure sustainable harvesting (NRET 2023; NRET 2024).

Although kangaroos typically affect plant regeneration on mainland Australia (Gordon *et al*. 2021; Morgan 2021), the Forester kangaroo (*Macropus giganteus tasmaniensis*) in Tasmania is now restricted to a few isolated populations in central and northeastern Tasmania (Lethbridge 2020). Unregulated hunting and habitat degradation reduced the Forester kangaroo in Tasmania to fewer than 15% of its pre-European range (Barker 1990; Tanner & Hocking 2001). Unlike more common macropod species such as Bennett’s wallaby and Tasmanian pademelon, Forester kangaroos are rarely detected during the statewide annual spotlight surveys, and the resulting low encounter rates provide insufficient data for robust population estimates. In recent years, the state government has undertaken aerial surveys with helicopters to improve the knowledge of species abundance and distribution of Forester kangaroos and to better inform future management decisions (Lethbridge 2020).

Drones have become an important tool in ecological research, particularly for wildlife monitoring (Corcoran 2019; Gazagne *et al*. 2023; Lappin *et al*. 2024). Unlike traditional ground-based methods such as spotlight surveys, drones can access remote or challenging habitats and can survey areas that are difficult, dangerous, or impossible to reach on foot (Chabot & Bird 2015; Povlsen *et al*. 2023). This capability is particularly relevant for macropod monitoring in Tasmania, where spotlight surveys remain the primary statewide detection method. Although widely used, spotlight surveys are labour intensive, subject to observer bias (Wayne *et al*. 2005; Sunde & Jessen 2013), and often yield low detection rates, especially in densely vegetated areas (Scott *et al*. 2005). Similarly, aerial surveys using helicopters or fixed-wing aircraft are costly (Clancy 1999; Barnas *et al*. 2018), logistically complex (Ulhaq *et al*. 2021), and pose significant safety risks; such operations have historically been a leading cause of death among field ecologists (Watts *et al*. 2010; Linchant *et al*. 2015).

Drone surveys address many of these limitations and offer several key advantages. They produce permanent, verifiable visual records that can be reviewed by multiple observers, reducing bias and allowing for quality control. This contrasts with traditional methods, where observations are typically recorded in real time without visual documentation. Drones also provide a safer and more cost-effective alternative to crewed aerial surveys, while offering greater repeatability in data collection (Linchant *et al*. 2015). When combined with automated detection techniques utilising Convolutional Neural Networks (CNNs), drone imagery can be processed efficiently at scale, enabling rapid analysis and reporting (Dujon *et al*. 2021; Lenzi *et al*. 2023). CNNs have been increasingly adopted in wildlife research over the past decade, with applications ranging from identifying species in camera trap images (Nguyen *et al*. 2017) to detecting marine megafauna in aerial surveys (Gray *et al*. 2019; Mannocci *et al*. 2022). These advances highlight the potential of deep learning to overcome long-standing challenges in ecological monitoring. Generalised modelling frameworks like `DeepForest` have been developed for object detection in aerial imagery, facilitating applications such as tree (Weinstein *et al*. 2020) and bird detection (Weinstein *et al*. 2021) in ecological studies.

In this study, transfer learning was applied to train a model for detecting macropods in drone imagery. Transfer learning repurposes a model trained on one task for another related task, enabling robust results even when training data are limited (Weinstein 2018; Gupta, Pathak & Kumar 2022). Most pre-built models are initially trained on very large datasets, such as ImageNet (Ribani & Marengoni 2019; Gupta, Pathak & Kumar 2022), which contain millions of labelled images of diverse objects. These models learn general visual features—such as edges, textures, and shapes—that can then be adapted to detect new targets, including wildlife in aerial imagery. Developing machine learning models from scratch typically requires extensive datasets and programming expertise. In contrast, transfer learning leverages existing knowledge from a pretrained model as a foundation, reducing both the data requirements and computational burden. Depending on the dataset size and complexity, either the entire model can be fine-tuned or only the final layers retrained, which allows the model to specialise for the new task without losing the general visual features learned during pretraining (Gupta, Pathak & Kumar 2022). This approach not only saves time and resources, but can also enhance model accuracy, particularly in data-scarce ecological contexts (Weinstein 2018). The aim of this study is to develop an automated detection technique, using transfer learning, to process drone imagery of macropods more efficiently. This study also investigated how environmental factors, particularly solar altitude angle, might influence the detectability of macropods across different times of day and habitats.

## 2. Methods

### 2.1 Study sites

The study took place at two study sites—Cockatoo Hills and Narawntapu National Park—both located in Tasmania, Australia. Cockatoo Hills (Lat: –42.23248, Lon: 146.47884) is situated in Tasmanian Central Highlands. Previously a private farmland, it is now a feeding ground for marsupials and other native species such as the bare-nosed wombats (*Vombatus ursinus*). The area is surrounded by dry sclerophyll forests (Teo *et al*. 2024). Narawntapu National Park (Lat: –41.149141, Lon: 146.603217) on Tasmania’s central north coast. The park has a vast landscape of coastal heathlands and open grasslands, which provides a suitable habitat for marsupial grazing (Martin *et al*. 2018; Teo *et al*. 2024).

### 2.2 Drones and flight operations

Two quadcopters, a DJI Phantom 4 Pro (DJI, Shenzhen, China) and a DJI Mavic 3 Thermal were used in this study. Both drone models are equipped with a visual (RGB) camera. The camera on the Phantom 4 Pro has a resolution of 5,472 × 3,648 pixels (sensor: 1’’ CMOS, effective pixels: 20M) and a maximum flight time of approximately 30 minutes without additional payload. The Mavic 3T has a maximum image resolution of 8,000 × 6,000 pixels (sensor: 1/2’’ CMOS, effective pixels: 48 MP). It has a maximum flight time of 45 minutes with no additional payload. Two different drone models were used, reflecting routine technological upgrades that occurred during the study. Initial checks confirmed no substantial differences in image quality or resolution between the DJI Phantom 4 Pro and DJI Mavic 3 Thermal under field conditions.

Drone surveys were conducted on clear, sunny days with low wind speed (<20 km/h) to optimise image quality. The launch site was located at least 150 metres from the nearest animal to minimise disturbance. Upon taking off, the drone was ascended to 40 metres above ground level (AGL), based on the species-specific protocol established in a previous study (Teo *et al*. 2024). This altitude balanced image resolution with minimal behavioural disturbance. All flight missions were pre-programmed using DJI Ground Station Pro, operating in a standard lawnmower (grid) pattern to ensure systematic coverage (Figure 1). Forward and side overlap were set to 80% to allow for robust image stitching and to reduce the risk of missing individuals due to movement or obstruction. The gimbal was set to –90° for consistent top-down imagery. The drone was flown at 5 ms⁻¹ with an appropriate shutter speed to minimise motion blur in the imagery. Missions were conducted across a range of times during daylight hours, from shortly after sunrise until before sunset, to intentionally sample under varying solar angles and lighting conditions, while also avoiding harsh shadows and low-light environments that could compromise image quality.

**Figure 1.**
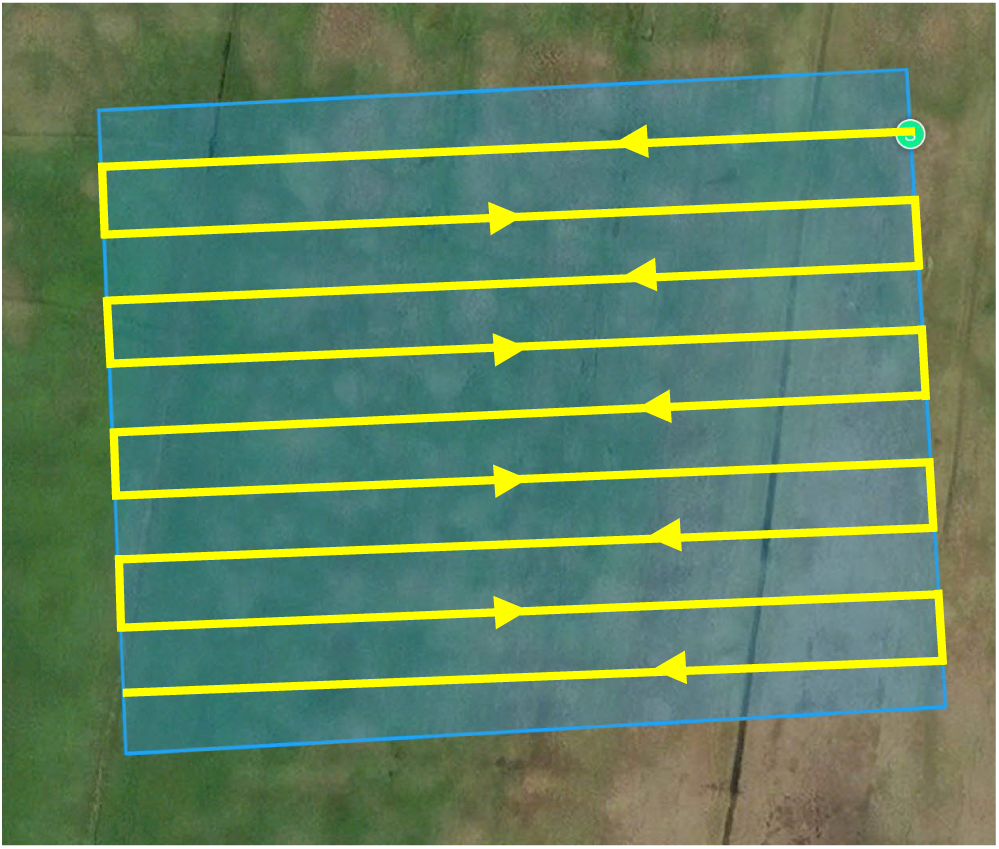
An example of a lawnmower pattern flight path.

### 2.3 Preparation of data

Drone imagery from both study sites was manually annotated with the software *labelImg* (https://pypi.org/project/labelImg/) (Figure 2). Bennett’s wallabies and Forester kangaroos were collectively labelled as ‘macropod’ to create a unified detection class, as distinguishing between the two species from aerial imagery is challenging and the available dataset was insufficient to support reliable species-level classification. Obscured animals (e.g., in deep shadows) were excluded from annotations to avoid compromising model accuracy. While this approach prioritised clean input data, it is noted that model performance may be slightly overestimated under field conditions, where partial visibility is more common. Individual drone images from both study sites were systematically cropped into 500 × 500-pixel windows, with each crop containing at least one annotated bounding box. Image sections without bounding boxes were excluded from training to streamline the dataset.

**Figure 2.**
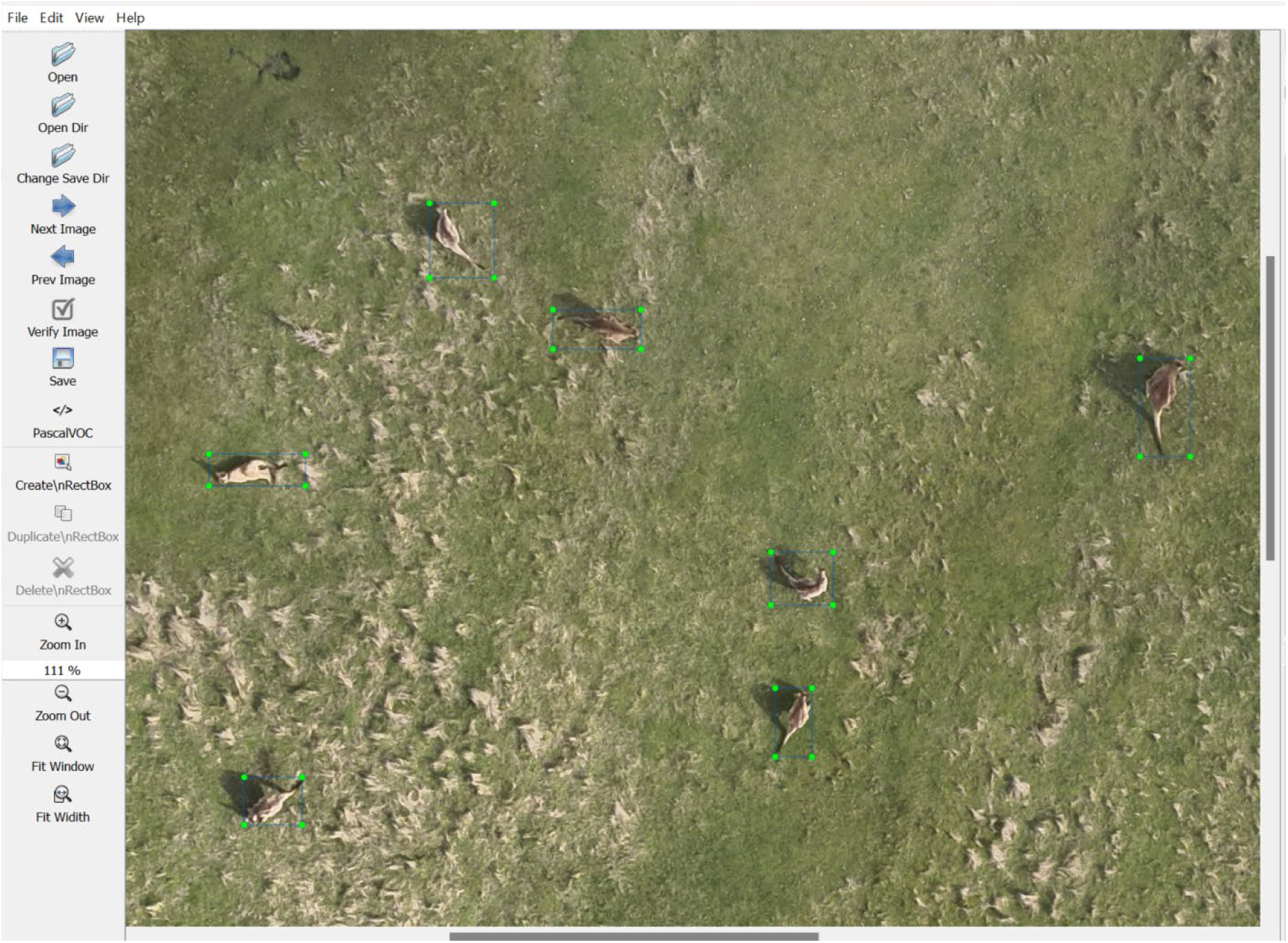
A screenshot of manual annotation using the *labelImg* software, showing macropods outlined with bounding boxes (in green) as part of the training dataset preparation.

The resulting datasets consisted of 4,129 cropped images from Cockatoo Hills and 8,385 cropped images from Narawntapu National Park. In addition to these two site-specific datasets, a third dataset was also created by merging them, resulting in a combined set of 12,514 cropped images. These three separate datasets were used to develop two ‘local models’ (trained exclusively on either Cockatoo Hills or Narawntapu National Park data) and one ‘joint model’ (trained on the combined dataset from both study sites). The joint model was trained from scratch using the merged data, without prior weighting or fine-tuning from the local models.

### 2.4 Architecture of a Convolutional Neural Network (CNN)

A Convolutional Neural Network (CNN) is a deep learning model widely used for image and video analysis, typically comprising convolutional, pooling, and fully connected layers (Gupta, Pathak & Kumar 2022). In this study, a CNN-based architecture was applied to extract spatial features from 500 × 500-pixel RGB drone images of macropods (Figure 3). These features were passed through a Region Proposal Network (RPN) and classification layers to detect and localise animals within the imagery. This structure follows the Faster R-CNN architecture (Ren *et al*. 2017), a two-stage object detection model in which the first stage proposes candidate bounding boxes and the second classifies and refines them. This architecture was implemented through the `DeepForest` framework (Weinstein *et al*. 2020), which was adapted to the drone imagery using transfer learning, as described in the following section.

**Figure 3.**
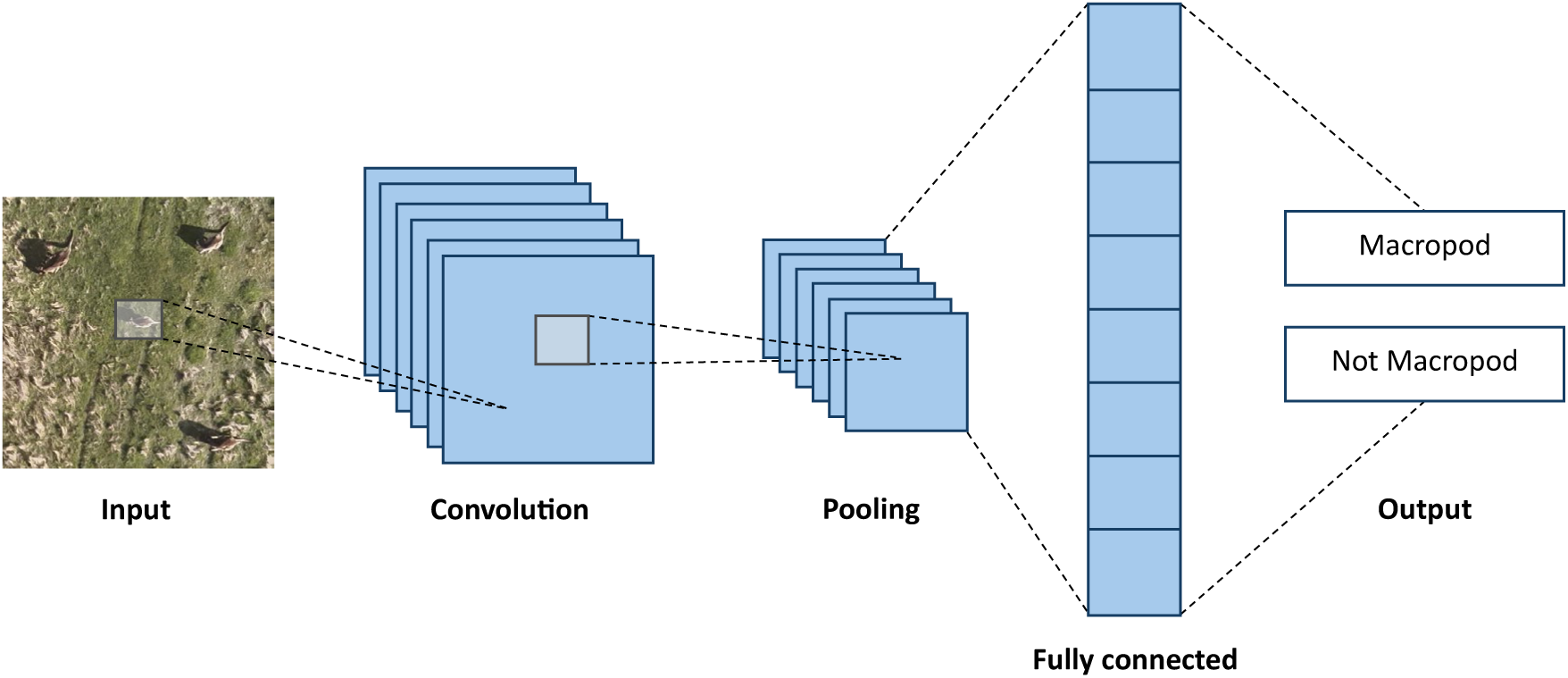
An illustration that demonstrates a simple CNN architecture which includes a convolutional layer, a pooling layer, and a fully connected layer.

### 2.5 Model training through transfer learning

`DeepForest`, a deep learning object detection model originally trained on tree crowns (Weinstein *et al*. 2020), was used and adapted via transfer learning for macropod detection. `DeepForest` is built on the Faster R-CNN architecture implemented in PyTorch’s `torchvision` module. `DeepForest` is designed for training and predicting ecological objects in airborne imagery (Weinstein *et al*. 2020). The `DeepForest` architecture provides a single-class object detection (tree crowns) model that can be extended to new object types through transfer learning. Unlike most wildlife detection models, which are typically trained on ground-based camera trap imagery, `DeepForest` was specifically trained on RGB aerial imagery. This makes it particularly well suited for adaptation to drone-based macropod detection, as it has already learned to detect natural, irregular shapes (tree crowns) in cluttered, vegetated environments from a top-down perspective. These shared visual and environmental features reduce the domain shift between the original and target tasks, improving the transferability of the pre-trained model to macropod detection.

The base model from the `DeepForest` package was initialised, and its pre-trained weights and biases were loaded into the training dataset. All three models (two local models and one joint model) were trained on a machine equipped with an NVIDIA Quadro RTX 5000 GPU with 16 gigabytes (GB) of memory. Fine-tuning was conducted by adjusting a few key hyperparameter values, including the number of training epochs, batch size, and image size based on dataset size and hardware constraints to optimise performance. All models were trained on 500×500-pixel images, using a batch size of 20 samples per iteration. Training was performed across 1 to 50 epochs, evaluated at 5-epoch intervals, to monitor model performance and identify the point of diminishing returns. This incremental approach was adopted, rather than training for 50 epochs outright, to assess learning progression and mitigate overfitting. Five-fold cross-validation was applied for all models. For the local models, folds were created separately within each site-specific dataset to evaluate performance on site-representative data. For the joint model, images from both sites were combined and randomly shuffled across folds to ensure generalisation across both environments.

### 2.6 Performance metrics

The models were assessed using the widely used performance metrics, which include intersection over union (IoU), box precision, box recall, and F1 score. *IOU* is calculated by dividing the area of overlap between the predicted bounding box and the ground truth bounding box by the total area covered by both boxes combined (i.e., the area of their union) as shown in Equation (1).

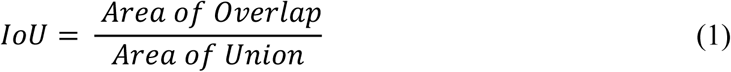

An *IOU* threshold of 0.4 was used to determine whether a predicted bounding box qualified as a true positive. Predictions with an *IOU* below this threshold were considered false positives, and unmatched ground truth boxes were treated as false negatives. This threshold is slightly lower than the commonly used 0.5 in general object detection benchmarks (Everingham *et al*. 2009) and was chosen to accommodate minor spatial offsets between predictions and annotations, which are common in aerial imagery due to animal movement, variation in image resolution, and slight human annotation inconsistencies.

*Box precision* measures the proportion of predicted bounding boxes that correctly overlap a ground truth box (Weinstein *et al*. 2020) as defined in Equation (2). A high box precision indicates that the model is accurately identifying the locations of objects in the image, with minimal false positives. A high box precision reflects the model’s ability to correctly place bounding boxes around the relevant objects with a high level of accuracy.

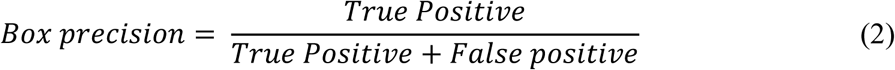

*Box recall* is the proportion of actual objects in the image that have been correctly detected by the model as shown in Equation (3), with a true positive defined as a predicted box overlapping the ground truth box according to the *IOU* threshold. A high box recall indicates that the model is effective at finding and localising the objects, with very few being missed (i.e., minimal false negatives).

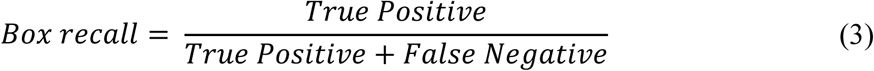

The *F*_1_ score is a measure of a model’s balance between precision and recall. The *F*_1_ score ranges from 0 to 1, where 1 indicates perfect precision and recall.

The *F*_1_ score is defined by Equation (4):

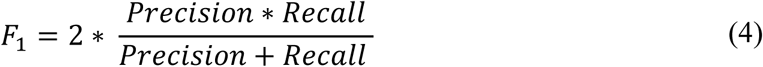

### 2.7 Orthomosaic Processing

To apply the trained model to field data, eight sets of drone images (four from Cockatoo Hills and four from Narawntapu National Park) were captured and processed using Agisoft Metashape Professional to create orthomosaics. Although the drone images used to generate the orthomosaics were collected with 80% forward and side overlap to ensure seamless stitching, the final orthomosaics were split into non-overlapping 500×500-pixel tiles for model inference. No additional overlap or post-processing was applied to the tiles during detection.

The resulting tile sets were then evaluated with the three trained models (two local models and one joint model) with the highest recall, as described in Section 3.1. Each model was applied to detect macropods, with the local models applied to their respective sites (Cockatoo Hills and Narawntapu National Park). For consistency, tile sets from these sites are hereafter referred to as ‘CH’ and ‘NNP’, respectively. The joint model was applied to both sites. Predicted tiles were then imported into ENVI (version 6.0) software for evaluation, where manual review was conducted to assess false positives and false negatives. In total, 16 tile sets (eight at each site) were analysed, with local and joint model outputs compared.

Solar altitude angles for each tile set were extracted from the Geoscience Australia azimuth calculator (Geoscience Australia 2025) based on the time and location of each flight. This variable was included to assess whether lighting conditions influenced detection performance.

The solar altitude angle refers to the angle between the sun’s position and the horizontal plane of the Earth’s surface as seen from an observer’s viewpoint (local horizon), with its maximum value occurring at noon (Ahmad *et al*. 2013). Solar altitude angle can affect the overall quality of drone imagery by affecting the length and direction of shadows cast on the ground, which can obscure details and distort features. As such, conducting drone surveys near midday, when the solar altitude angle approaches 90°, is recommended to minimise shadowing (Puttock *et al*. 2015). In this study, the effect of solar altitude angle on macropod detection performance across different times of day and habitat types was tested by comparing F1 scores across the corresponding solar altitude angles for each flight.

## 3. Results

### 3.1 Model performance

The performance metrics for both the local and joint models were evaluated at each epoch during training (Figure 4). Model selection was based on the highest recall score, as this metric prioritises the detection of all macropods in the imagery, which is crucial for accurate population monitoring.

**Figure 4.**
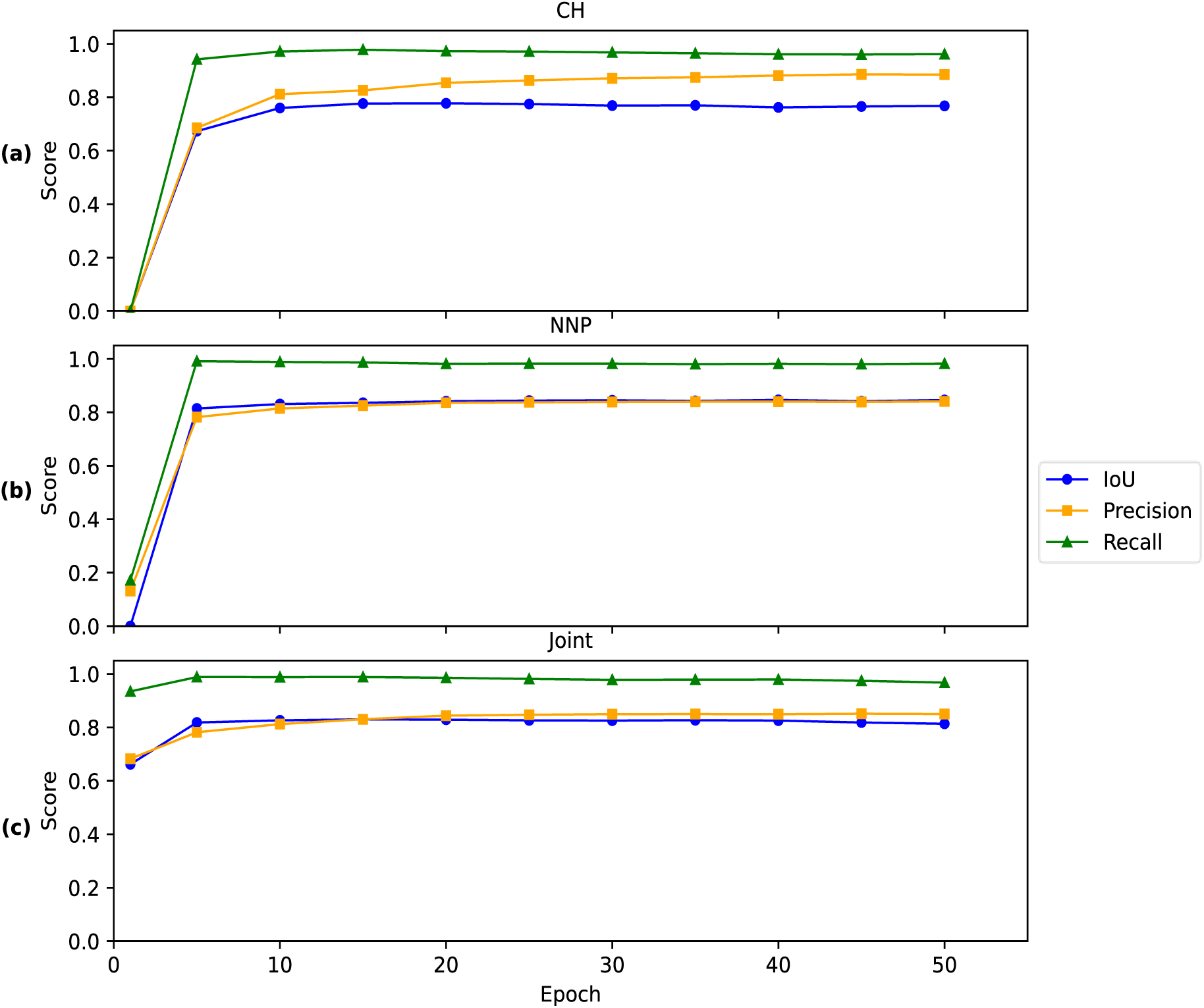
Performance metrics for macropod detection models across training epochs. Graph panels show model performance for (a) Cockatoo Hills local model (CH), (b) Narawntapu National Park local model (NNP), and (c) joint model (Joint).

The Cockatoo Hills model achieved optimal recall (0.978) at 15 epochs, with a corresponding precision of 0.826 (Figure 4a). The Narawntapu model reached 0.991 at 5 epochs, with a precision of 0.782 (Figure 4b). The joint model attained 0.988 at 15 epochs, with a precision of 0.831 (Figure 4c). All models demonstrated the characteristic learning curve of convolutional neural networks, with rapid performance improvement during early epochs followed by stabilisation. The final model for each configuration was selected based on maximising recall, with consideration of acceptable trade-offs in precision.

### 3.2 Evaluation

The F1 scores for all 16 sets—eight from Cockatoo Hills (CH) and eight from Narawntapu National Park (NNP)—were calculated based on precision and recall metrics (Table 1 and Table 2). Across all datasets, the joint model applied to dataset NNP 4 yielded the highest F1 score of 0.83, with a precision of 0.93 and recall of 0.76 (Table 2). The corresponding confusion matrix (Figure 5) shows 50 true negatives and 37 true positives, with only 3 false positives and 12 false negatives, highlighting the joint model’s strong overall performance under these conditions.

**Figure 5.**
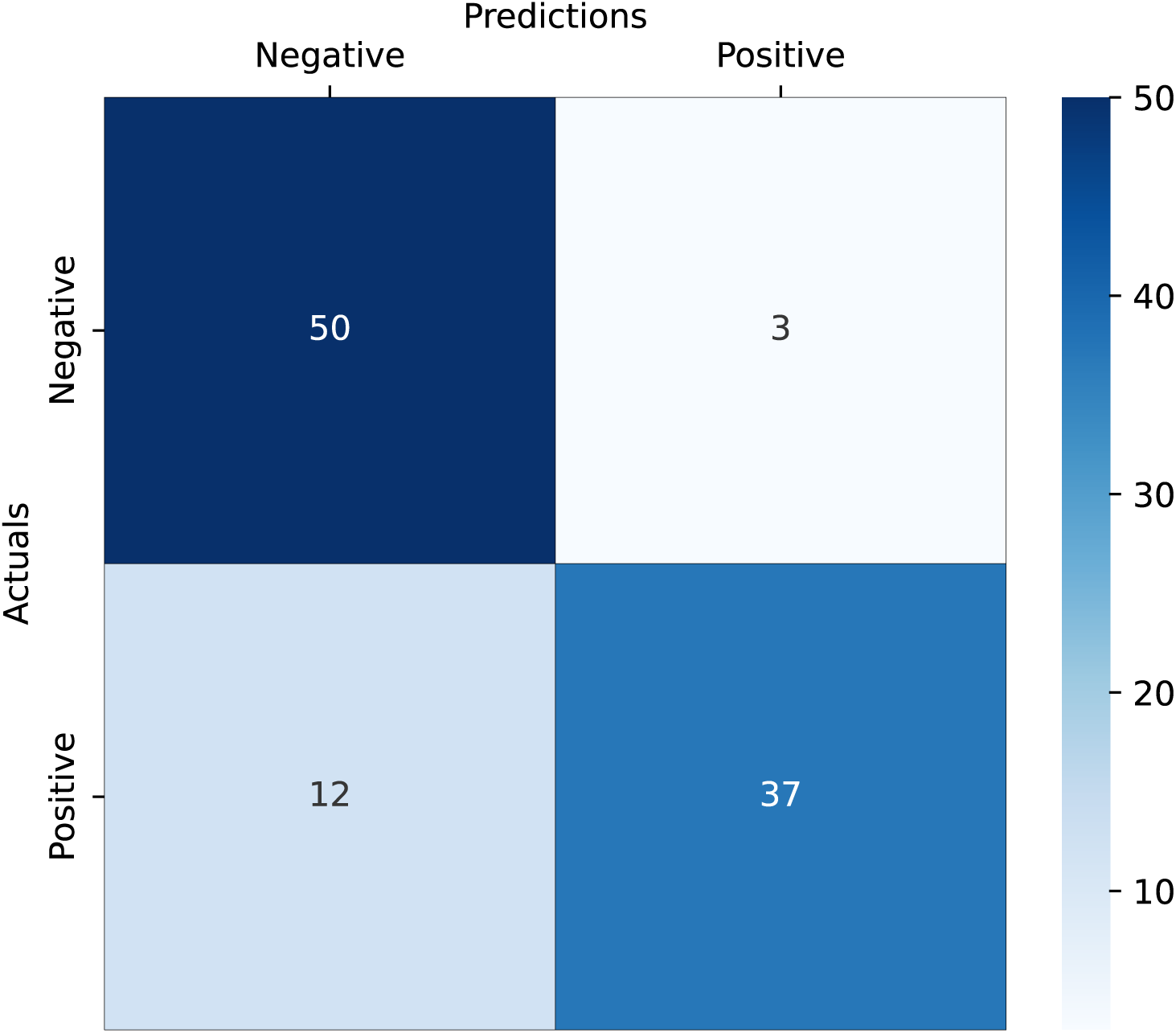
Confusion matrix of Dataset NNP 4 using the joint model (highest F1 score among all datasets). Colour intensity corresponds to the magnitude of each cell count. The matrix shows correct and incorrect predictions (negative vs. positive classes): 50 true negatives, 37 true positives, 3 false positives, and 12 false negatives.

**Table 1.**
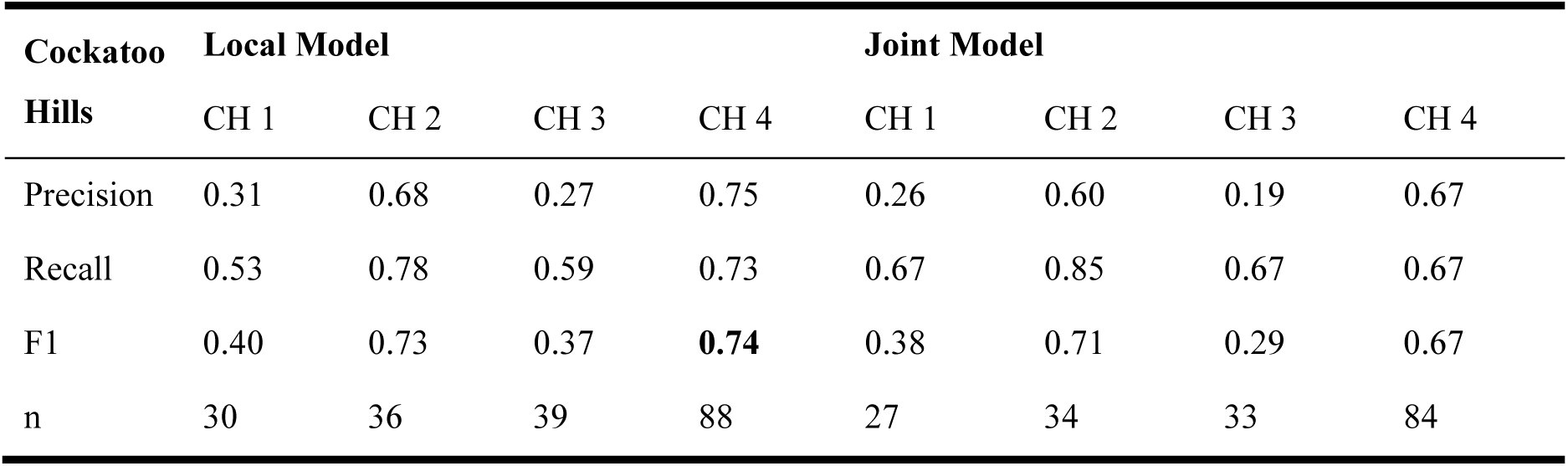
Precision, recall, and F1 scores for macropod detection using the local and joint models across four datasets (CH 1 to CH 4) at Cockatoo Hills. The sample size (n) represents the number of ground truth macropods present in each dataset. The highest F1 score (0.74) was achieved with the local model on dataset CH 4.

| Cockatoo Hills | Local Model |  |  |  | Joint Model |  |  |  |
| --- | --- | --- | --- | --- | --- | --- | --- | --- |
|  | CH 1 | CH 2 | CH 3 | CH 4 | CH 1 | CH 2 | CH 3 | CH 4 |
| Precision | 0.31 | 0.68 | 0.27 | 0.75 | 0.26 | 0.60 | 0.19 | 0.67 |
| Recall | 0.53 | 0.78 | 0.59 | 0.73 | 0.67 | 0.85 | 0.67 | 0.67 |
| F1 | 0.40 | 0.73 | 0.37 | **0.74** | 0.38 | 0.71 | 0.29 | 0.67 |
| n | 30 | 36 | 39 | 88 | 27 | 34 | 33 | 84 |

**Table 2.** Precision, recall, and F1 scores for macropod detection using the local and joint models across four datasets (NNP 1 to NNP 4) at Narawntapu National Park. The sample size (n) represents the number of ground truth macropods present in each dataset. The highest F1 score (0.83) was achieved with the joint model on dataset NNP 4.

| Narawntapu National Park | Local Model |  |  |  | Joint Model |  |  |  |
| --- | --- | --- | --- | --- | --- | --- | --- | --- |
|  | NNP 1 | NNP 2 | NNP 3 | NNP 4 | NNP 1 | NNP 2 | NNP 3 | NNP 4 |
| Precision | 0.77 | 0.49 | 0.25 | 0.68 | 0.79 | 0.59 | 0.17 | 0.93 |
| Recall | 0.80 | 0.79 | 0.57 | 0.79 | 0.83 | 0.70 | 0.40 | 0.76 |
| F1 | 0.78 | 0.60 | 0.35 | 0.73 | 0.81 | 0.64 | 0.24 | **0.83** |
| n | 25 | 24 | 7 | 52 | 23 | 23 | 5 | 49 |

In general, NNP datasets outperformed CH datasets across all metrics, particularly under the joint model. At CH, the highest F1 score was 0.74, achieved by the local model on CH 4 (Table 1). Performance at CH was more variable, and precision tended to be lower than at NNP, likely due to the denser vegetation and more complex background.

To investigate environmental influences on detection performance, F1 scores were compared with the solar altitude angle at the time of each flight (Figure 6). A clear declining trend was observed: F1 scores for both sites decreased as solar altitude increased, with the most accurate results occurring when the solar angle was below 30°. This pattern held for both local and joint models. For example, the NNP joint model achieved F1 scores of 0.81 and 0.83 at lower solar angles (∼16-28°), but dropped to 0.24 at the highest angle (∼49°). A similar trend was observed in CH datasets, where F1 scores declined from 0.74 to 0.29 as solar angle increased.

**Figure 6.**
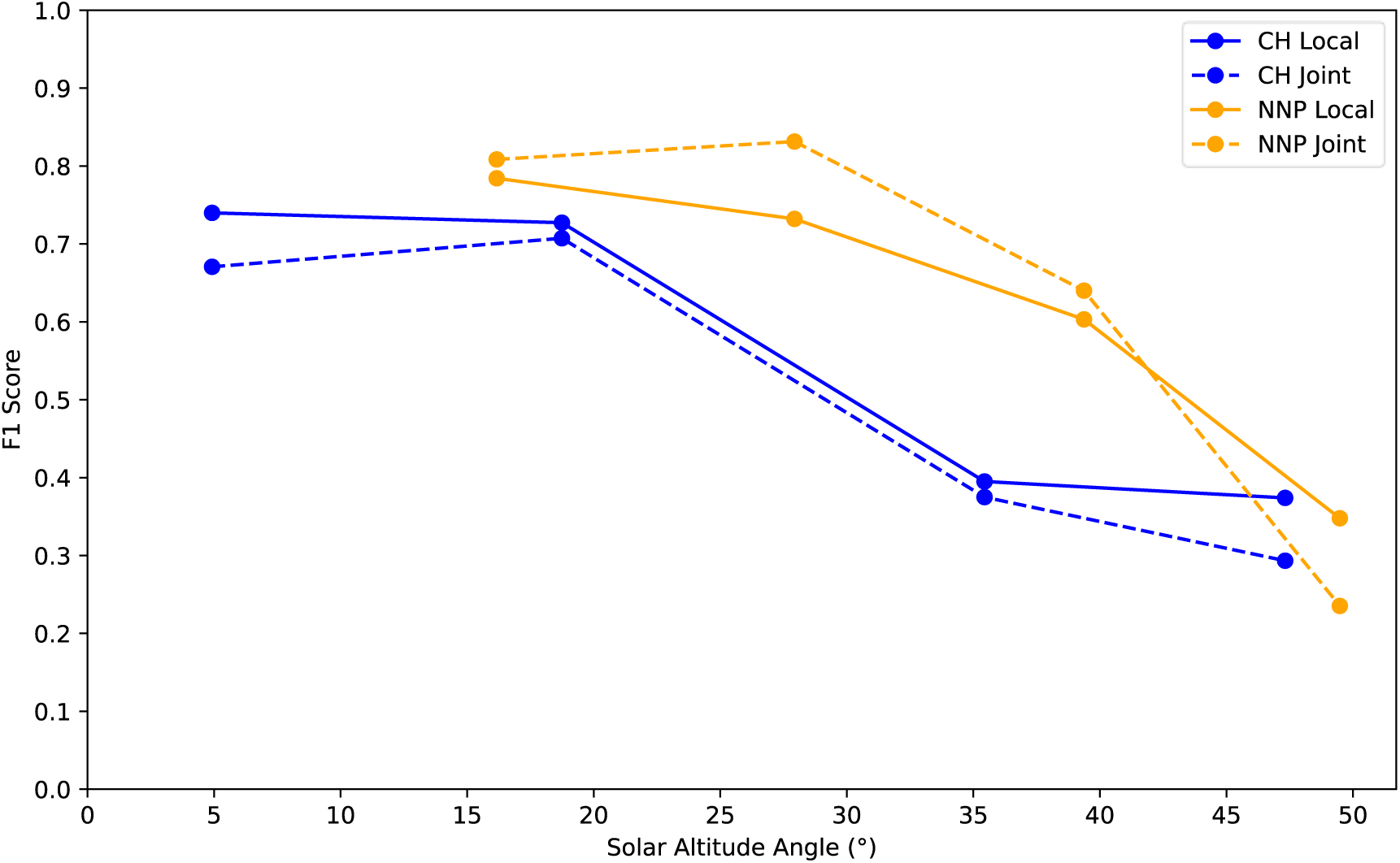
F1 scores of local and joint models for macropod detection at Cockatoo Hills (CH) and Narawntapu National Park (NNP) plotted against solar altitude angle. Both local and joint models exhibit declining F1 scores as solar altitude increases, with the highest performance below 30°. Solar altitude angles were obtained from Geoscience Australia’s azimuth calculator (Geoscience Australia 2025).

Figure 7 illustrates how solar altitude varied over the survey days, peaking around noon (∼50°) and decreasing in the early morning and late afternoon.

**Figure 7.**
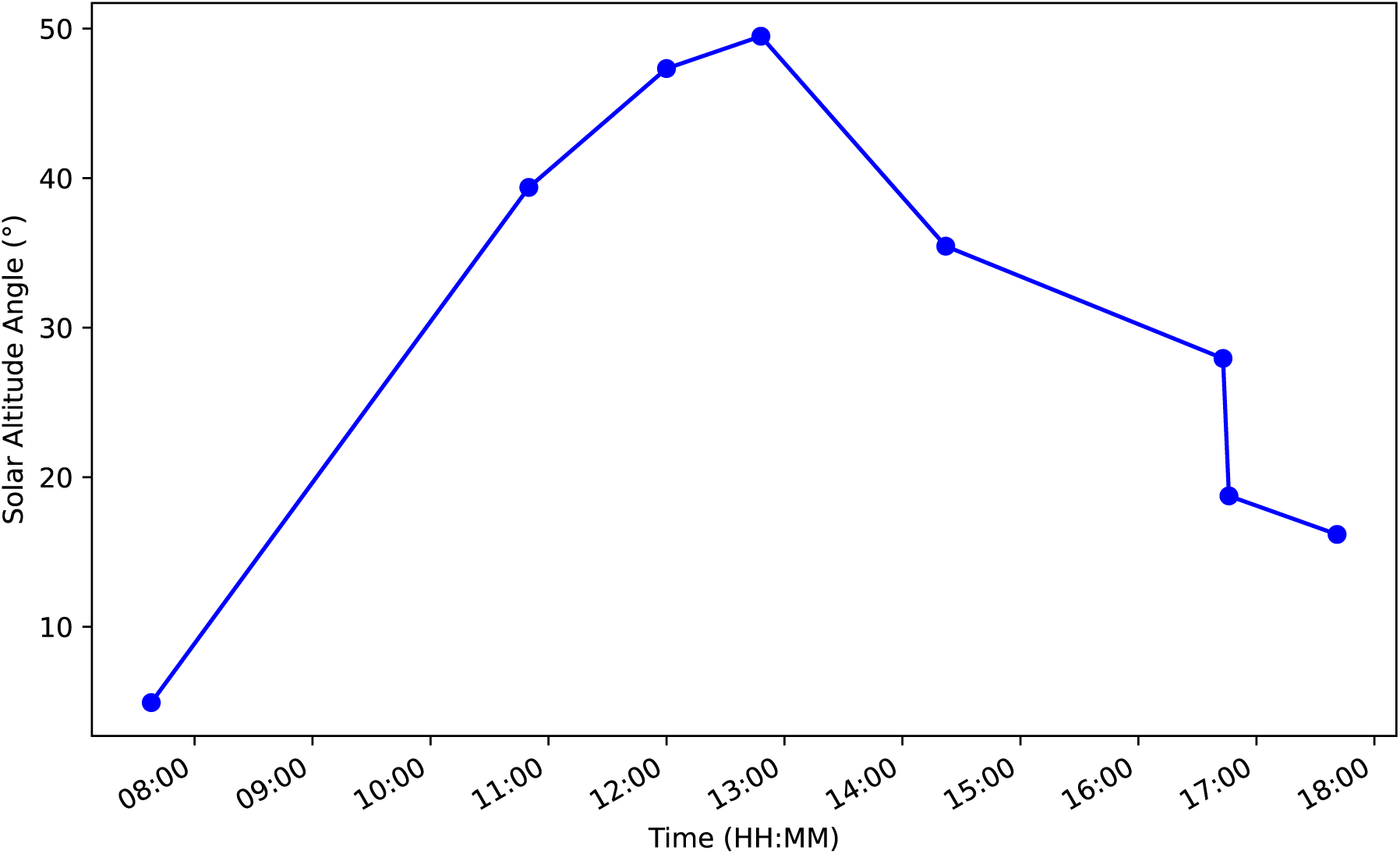
Solar altitude angles recorded during drone surveys conducted on different days in autumn and spring, where daylight hours were similar (approximately 12 hours). The solar altitude angle peaks around noon (between 12:00 and 13:00), reaching approximately 50°, and decreases toward the late afternoon. The lowest angles are observed in the early morning and late evening. Solar altitude angles were obtained from Geoscience Australia’s azimuth calculator (Geoscience Australia 2025).

## 4. Discussion

This study evaluated the performance of a transfer learning-based object detection model for macropod monitoring using drone imagery across two contrasting Tasmanian habitats. The CNN model, adapted from `DeepForest`, achieved strong detection results at both sites, with F1 scores exceeding 0.7 in several datasets, demonstrating good generalisability despite habitat differences. Key findings showed that detection performance was highest at lower solar altitude angles (<30°), particularly in vegetated areas, and that recall-prioritised model selection offered a balanced trade-off between missed detections and false positives. Limitations such as reduced visibility in dense cover and the absence of negative samples were identified as areas for future model improvement.

### 4.1 Field surveys

In this study, drones were pre-programmed to operate in a lawnmower pattern (also called a grid pattern), which allows for even coverage of the survey area in a structured and systematic manner, thereby minimising data collection gaps. This flight pattern captures overlapping images from adjacent flight paths, which is important for post-processing tasks such as creating orthomosaics or digital elevation models. The overlap ensures that enough data points are available for accurate image stitching. However, animals in overlapping areas may appear in multiple images, leading to potential double counting (Brack *et al*. 2018). For instance, a macropod detected in one image may move to an adjacent area while the drone proceeds along its flight path, resulting in the same individual being recorded again in subsequent images (Brack *et al*. 2018). Furthermore, if the drone captures images at fixed intervals while hovering, rather than capturing continuously in flight, animals could relocate between those intervals, increasing the likelihood of double counting (Brack *et al*. 2023). This highlights an ongoing challenge in drone-based wildlife surveys, where the need for high image overlap must be balanced with the potential for movement-based duplication. To mitigate this, future studies could explore flight scheduling strategies that minimise the time gap between successive passes—such as reducing transect spacing or using faster flight speeds—while still maintaining sufficient overlap for post-processing.

The transfer learning-based model developed in this study achieved robust macropod detection across both open and vegetated habitats, demonstrating strong performance in environments that typically challenge ground-based methods. In contrast, traditional spotlight surveys are often constrained by poor visibility, particularly in dense vegetation, and are prone to observer bias and variability due to environmental conditions (Fletcher, Moller & Clapperton 1999; Wayne *et al*. 2005; Sunde & Jessen 2013). While drone surveys are also influenced by environmental conditions, these can be minimised through careful planning (e.g. scheduling flights under favourable light and weather conditions). Moreover, a consistent top-down perspective and systematic coverage were provided through the use of pre-programmed flight paths. This standardised approach reduced spatial bias and improved detection reliability, even in more complex terrain (Preston *et al*. 2021; Povlsen *et al*. 2023). However, a direct comparison with ground-based or crewed aerial methods on the same populations would be required to confirm this more conclusively.

When carrying out drone surveys during daylight hours, solar altitude angle—rather than simply the time of day—has a significant impact on image quality. The results indicated that detection performance was higher at lower solar altitude angles, with all F1 scores exceeding 0.6 when the angle fell below 30° (i.e., before 9:30 AM and after 4:30 PM). This finding contrasts with the recommendations of Puttock *et al*. (2015), who suggested flying within a few hours of midday to minimise shadowing. In this study, however, midday conditions produced harsh, high-contrast shadows, particularly at Cockatoo Hills.

This could be attributed to the complexity of the habitat in the study area. At Narawntapu National Park, where the study sites were primarily open grasslands with minimal vegetation, shadowing had minimal impact on image quality. The absence of dense cover allowed for more consistent visibility of macropods throughout the day (Figure 8c & d). At Cockatoo Hills, where vegetation is denser, higher solar angles (>30°) produced harsh shadows that obscured details, reducing the effectiveness of detection (Figure 8a). Better performance at angles below 30° likely reflected softer, more diffuse lighting that diminished shadow contrast, thereby enhancing detection in the denser vegetative environment (Figure 8b). These findings suggest that optimal survey timing should be adjusted according to habitat structure, and that lower solar angles may improve detectability in denser environments. In such conditions, thermal imaging, which does not rely on visible light, may provide a practical means of reducing shadow-related detection issues, especially in dense or heterogeneous environments such as Cockatoo Hills.

**Figure 8.**
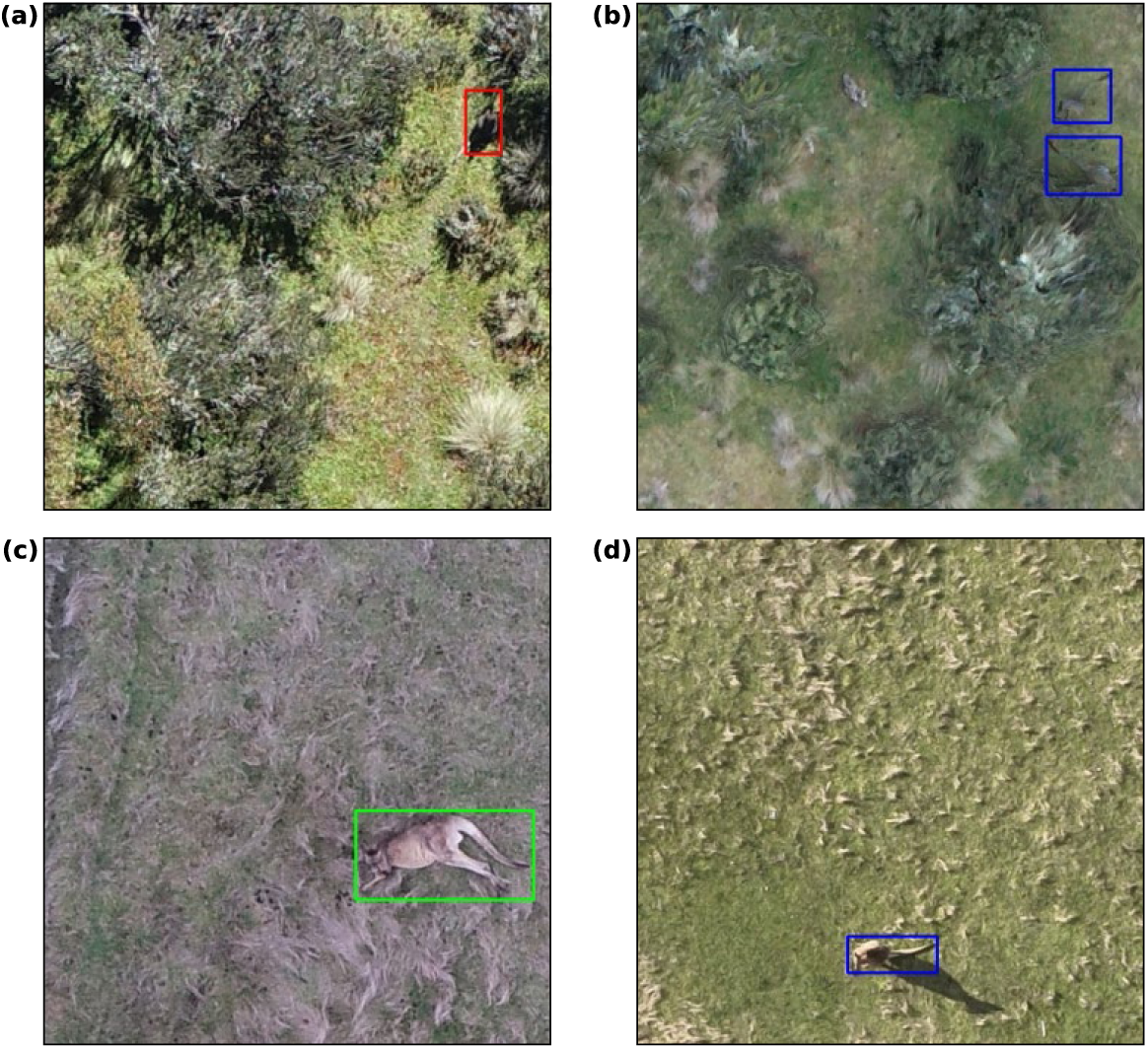
Effect of habitat complexity and solar altitude on drone-based macropod detection. Bounding boxes are color-coded based on the model’s confidence: green (>90%), blue (51-90%), and red (21-50%). Drone imagery shows macropods (highlighted in bounding boxes) in complex habitat at Cockatoo Hills captured at 12:00 PM (a) and 7:38 AM (b), and in the less complex habitat at Narawntapu National Park at 12:48 PM (c) and 4:43 PM (d).

Since both study sites were surveyed during autumn and spring, daylight hours were similar (approximately 12 hours), resulting in comparable solar altitude angles for the respective times of day. However, time windows shift across seasons. For instance, in winter, the sun sets earlier, meaning that a solar altitude angle at 4:30 PM in spring would not be the same in winter. The results suggest that solar altitude angle, rather than time of day alone, is a more reliable predictor of image quality and detection performance, particularly in structurally complex habitats. This finding builds on and refines previous assumptions that time-based scheduling is sufficient for planning drone flights. Accordingly, it is recommended that survey planning prioritise solar altitude thresholds over fixed times. Additionally, aligning flight operations with periods of peak species activity is important, especially for crepuscular or nocturnal animals such as macropods. Therefore, it is essential to carefully plan flight operations by considering solar altitude angles, target species’ activity patterns, habitat complexity, and available daylight hours to optimise the effectiveness of drone surveys.

While disturbance minimisation was not a primary aim of this study, flights were conducted using an established species-specific protocol (Teo *et al*. 2024) to minimise disturbance to the macropods. No overt behavioural responses were observed during drone operations. However, different species may respond to drone presence differently (Ditmer *et al*. 2015; Vas *et al*. 2015; Christiansen *et al*. 2020). To date, only one study has established drone disturbance thresholds for macropods: Brunton *et al*. (2019) recommended a flight altitude of 60-100 metres AGL for observing eastern grey kangaroos (*Macropus giganteus*), noting minimal behavioural response within this range. Further research on species-specific thresholds would nonetheless support the broader application of drone surveys in conservation and monitoring.

### 4.2 Image processing

The top-performing model from each dataset (two local, one joint) was selected by its highest recall, prioritising the detection of as many macropods as possible. Recall reflects the proportion of animals that were correctly identified out of the total present, which in population studies are important. Although choosing the best-performing model by recall may lead to slightly more false positives (e.g., misidentifying non-animal features as macropods), this trade-off was acceptable in this case. The selected models still achieved F1 scores close to the maximum for each site, indicating a good balance between recall and precision. For example, at Cockatoo Hills, the model with the highest recall had an F1 score of 0.71 compared to the maximum possible F1 of 0.74 (Table 1), and at Narawntapu National Park, the best recall model had an F1 score of 0.81 compared to the peak value of 0.83 (Table 2). These small differences suggest that prioritising recall did not significantly compromise overall model performance. It should be noted that even with high recall and F1 scores, uncertainty remains in population counts derived from drone surveys due to factors such as false negatives, variable image quality, and environmental influences on detectability. These uncertainties would result in relatively wide confidence intervals around estimated population sizes, meaning that only substantial changes in population numbers could be reliably detected over time. As with many wildlife survey methods, this limitation should be considered when applying drone-based detection for population trend assessments, and repeated surveys or complementary methods may be necessary to detect smaller or gradual changes.

However, model performance could still improve by reducing false positives and negatives, especially in complex habitats where the background and contrast between animals and their surroundings can complicate detection. The study site at Cockatoo Hills was characterised by dense vegetation, including shrubs and patches of vegetation, where macropods move between dry sclerophyll forests and dirt tracks. In contrast, the study site at Narawntapu National Park consisted of open grasslands, which were previously used as grazing land for livestock. The differences in habitat complexity between the two sites were reflected in the results, where the F1 score and recall for the Narawntapu National Park dataset were generally higher than those for Cockatoo Hills (Table 1 and Table 2). The dataset from Narawntapu National Park exhibited a higher contrast between the macropods and their surroundings, as the animals’ colour and shape stood out distinctly against the grassy background. Conversely, drone images from Cockatoo Hills showed lower contrast, with the macropods blending more into the dense vegetation, making detection more challenging.

Factors such as the animals’ body shape, size, and movement patterns, as well as variations in habitat between study sites, can influence the model’s performance (Norouzzadeh *et al*. 2018; Perz *et al*. 2023). Notably, image augmentation techniques were not applied, nor were negative samples (i.e. images without macropods), although both could potentially improve the model’s ability to generalise across a wider range of environmental conditions (Dujon *et al*. 2021). Negative samples are known to help reduce false positives by improving the model’s ability to distinguish target animals from background features. These approaches were not implemented here due to the limited dataset, where adding large numbers of negatives risked creating strong class imbalance and unstable training outcomes. Nonetheless, applying augmentation and negative samples represents a valuable direction for future research and could potentially enhance the model’s ability to generalise across a wider range of environmental conditions.

Another critical factor is the dataset size, as larger datasets typically result in more robust models. Increasing the dataset in future research would likely help reduce both false positives and false negatives, leading to more accurate detection outcomes. From a management perspective, false negatives (failing to detect animals that are present) may lead to underestimations of population size, which could delay intervention or mask overabundance issues in conservation areas or agricultural zones. Conversely, false positives (misclassifying background features as macropods) can inflate population estimates and potentially result in overly aggressive control measures (Kellenberger, Marcos & Tuia 2018). While this study prioritised recall to minimise missed detections, reducing both types of error remains critical for ensuring the reliability of drone-based surveys as a tool for evidence-based decision-making.

The ability of drones to repeatedly fly over the same area at different times of the day allows researchers to capture temporarily overlapping imagery, which helps reduce misidentifications by recognising static features that are not animals, as animals are dynamic and move across the landscape. Each drone image is geotagged, enabling researchers to pinpoint the exact location of an animal and reduce errors. For instance, if drone surveys are conducted at the same study site multiple times, an object or animal can be tracked across images. A stationary feature (e.g., a log that might be mistaken for a macropod in one image) can be ruled out if it remains in the same position across multiple images, whereas an animal would most likely have moved. Additionally, the images collected in the field provide valuable spatial information. Orthomosaics allow researchers to pinpoint exact locations of animals and link these to habitat features, such as proximity to edges or open areas. This spatial data enhances the understanding of the target species’ behaviour and habitat preferences, which is crucial for effective conservation management. Nevertheless, even when automated detection fails or proves less effective, the permanent data obtained from drone surveys (i.e., images or videos) remains valuable, as researchers can manually review the drone images and footage.

Finally, it is important to acknowledge constraints that may have limited model performance in this study. Daylight-based RGB imagery can hinder the detection of macropods sheltering under dense vegetation or otherwise obscured from view. While a species-specific flight protocol designed to minimise behavioural disturbance (Teo *et al*. 2024) was followed, this also meant avoiding low altitudes, which may have yielded more informative imagery. Future work could explore the use of thermal sensors to enhance detection under dense canopy cover or during crepuscular periods of peak animal activity. Ethical considerations, such as avoiding breeding seasons and minimising drone-wildlife interactions (e.g., with raptors), must also be factored into study design (Lyons *et al*. 2018), particularly when seeking to expand training datasets across different environments.

## 5. Conclusions

This study developed and evaluated an automated detection technique for identifying macropods in drone imagery using transfer learning. The CNN model, adapted from `DeepForest` and trained on annotated drone images from two Tasmanian sites, achieved strong detection performance across contrasting habitats. Models prioritising recall produced F1 scores close to the maximum, demonstrating their utility for minimising missed detections without greatly increasing false positives. These results confirm the feasibility of applying transfer learning to aerial wildlife monitoring, even with limited training data, and highlight the potential for deploying such models in routine macropod surveys. While performance varied with habitat complexity and solar altitude angle, the model’s adaptability suggests broad applicability across similar ecological contexts. Future research should expand training datasets, incorporate negative samples, and explore the use of thermal imagery to enhance detection under low-light or dense vegetation conditions. Although the primary aim of this study was to develop and evaluate automated detection, the outputs from these models could be combined with statistical approaches that account for imperfect detection—such as distance sampling or N-mixture models—to estimate population density and abundance, demonstrating the potential for applying automated detection in population-level monitoring.

## Notes

### Competing Interest Statement

The authors have declared no competing interest.

### Funding

- Australian Research Council: FL160100101

